# Cardiac enriched BAF chromatin remodeling complex subunit Baf60c regulates gene expression programs essential for heart development and function

**DOI:** 10.1101/184143

**Authors:** Xin Sun, Swetansu K. Hota, Yu-Qing Zhou, Stefanie Novak, Dario Miguel-Perez, Danos Christodoulou, Christine E. Seidman, J.G. Seidman, Carol C. Gregorio, R. Mark Henkelman, Janet Rossant, Benoit G. Bruneau

## Abstract

How gene networks controlling organ-specific properties are modulated by chromatin remodeling complexes is not well understood. *Baf60c* (*Smarcd3*) encodes a cardiac-enriched subunit of the SWI/SNF-like BAF chromatin complex. Its role throughout heart development is not fully understood. We show that constitutive loss of *Baf60c* leads to embryonic cardiac hypoplasia and pronounced cardiac dysfunction. Conditional deletion of *Baf60c* in cardiomyocytes results in postnatal dilated cardiomyopathy with impaired contractile function. *Baf60c* regulates a gene expression program that includes genes encoding contractile proteins, modulators of sarcomere function, and cardiac metabolic genes. Many of the genes deregulated in *Baf60c* null embryos are targets of the MEF2/SRF co-factor Myocardin (MYOCD). In a yeast two-hybrid screen we identify MYOCD as a BAF60c interacting factor; we show that BAF60c and MYOCD directly and functionally interact. We conclude that Baf60c is essential for coordinating a program of gene expression that regulates the fundamental functional properties of cardiomyocytes.

## Introduction

Transcription factor networks control cardiac morphogenesis and cell specification (Bruneau, 2013; Evans et al., 2010) including the coordinated regulation of genes encoding the proteins involved in sarcomere function (Creemers et al., 2006; Niu et al., 2008). While undergoing complex morphogenetic changes, the developing heart functions to support the embryonic circulation. The contractile function of the heart adapts quickly to the dramatic changes in circulation that occur after birth, and subsequently must adapt to fluctuating physiology and stress. The transcriptional regulation of cardiac gene expression continues during postnatal heart growth and cardiomyocyte maintenance (Huang et al., 2009; Oka et al., 2006).

Chromatin remodeling complexes are critical regulators of cardiac gene expression, in many cases modulating the activity of DNA-binding transcription factors (Chang and Bruneau, 2012). For example, histone deacetylases (HDACs) and Bromodomain-containing factors play important roles in cardiac gene regulation and remodeling, and have been proposed as potential therapeutic drug targets (Anand et al., 2013; McKinsey, 2012). BRG1/BRM-Associated Factor (BAF) complexes are ATP-dependent chromatin remodeling complexes related to the yeast SWI/SNF complex, and are indispensable for mammalian development (Hota and Bruneau, 2016). BAF complexes orchestrate many aspects of heart development, and genetically interact with cardiac transcription factors to finely modulate cardiac gene expression (Hang et al., 2010; Takeuchi et al., 2011). Combinatorial assembly of different polymorphic subunits can generate hundreds of potential BAF complexes, and offer precise control of developmental processes (Chang and Bruneau, 2012; Ho and Crabtree, 2010). BAF60c (also known as SMARCD3) is a polymorphic subunit of the BAF complex, which is expressed preferentially in the developing heart (Lickert et al., 2004). In vivo RNAi knockdown in mouse embryos suggested that *Baf60c* is essential for embryonic heart development (Lickert et al., 2004), and together with the cardiac transcription factors TBX5, NKX2-5 and GATA4, BAF60c can induce non-cardiac mesoderm to differentiate into cardiomyocytes (Lou et al., 2011; Takeuchi and Bruneau, 2009).

In this study, we examined the role of *Baf60c* in embryonic and postnatal heart development using a *Baf60c* conditional knockout mouse line. We show that *Baf60c* is essential for cardiac growth and cardiomyocyte function at several stages of embryonic development, by regulating broad networks of genes encoding proteins essential for function of the contractile apparatus. Many of the dysregulated genes are targets of the MEF2 co-factor MYOCD, and we identify MYOCD as a BAF60c-interacting protein. Our work shows that *Baf60c* serves as an important modulator of the fundamental program of gene expression essential for cardiac structure and function.

## Results

### Construction of *Baf60c* conditional knockout mouse line

*Baf60c* is expressed at E7.5 in the early cardiac precursors of the cardiac crescent, and its expression is maintained throughout development in the myocardium (Lickert et al., 2004). In order to understand the function of *Baf60c* at different developmental stages, we developed a conditional allele of *Baf60c* in the mouse. A targeting construct with a pair of loxP sites flanking exon 1-4 was introduced into embryonic stem (ES) cells (Fig 1A). Transgenic mice generated from the targeted ES cells (*Baf60c*^*flox*/+^) have normal phenotypes and life span and thus are treated as wild type. By crossing with pCAGGS-Cre mice which constitutively express Cre recombinase, exons 1-4 of *Baf60c* were deleted to generate *Baf60c*^+/-^ mice (Fig 1A). No obvious defects were observed in *Baf60c*^+/-^ mice. Homozygous null *Baf60c*^-/-^ embryos were recovered at E9.5 (Fig 1B), and by whole mount in-situ hybridization no *Baf60c* mRNA was detectable in *Baf60c*^-/-^embryos (Fig 1C).

**Fig 1.**
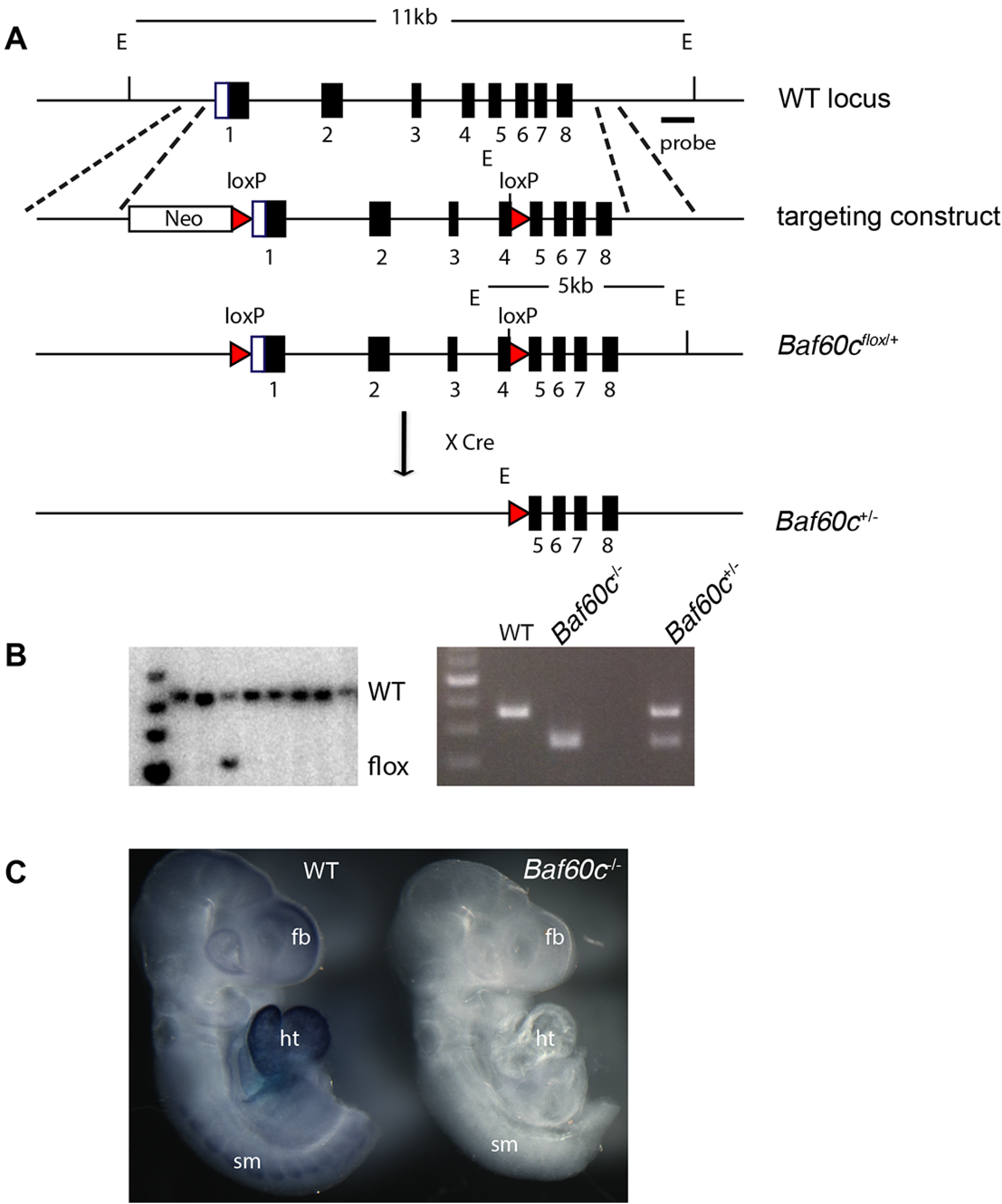
Construction of *Baf60c* knockout mouse line. A: Schematic representation of targeting DNA introduced into wild type (WT) *Baf60c* locus. Correctly targeted ES cells were identified with probe located outside of the homologous arm. Cre-mediated excision removed exon 1-4 and resulted in *Baf60c*^*+/-*^. B: Left: Southern blot of digested ES cell DNA with an external probe outside of the targeting DNA. WT and targeted band size are as described in A. Right: genotype PCR showing the band size difference of WT, *Baf60c*^*-/-*^ (KO) and heterozygous *Baf60c*^*+/-*^. C: Whole-mount in-situ hybridization using full-length Baf60c probe detected no signals in genotyped homozygous Baf60c^-/-^ embryos (n>3), indicating complete deletion. E: EcoRI; Tg: targeted; fb: forebrain; ht: heart; sm: somites.

### *Baf60c* deletion results in a hypoplastic heart and embryonic demise

*Baf60c*^-/-^ embryos were recovered alive and with roughly normal morphology at different stages of timed pregnancy until E12.5-E14.5. At E14.5, most *Baf60c*^-/-^ embryos were dead, with broad regions of hemorrhage. Backcrossing into C57Bl/6 for 10 generations led to a more consistent phenotype, with survival only until E12.5-13.5. To determine the cause of embryonic death, and to identify potential cardiac phenotypes, *Baf60c*^-/*-*^embryos were harvested for histological analysis. Optical projection tomography showed that mixed background E12.5 *Baf60c*^-/-^ embryonic hearts had dilated inner chambers and underdeveloped interventricular septa (Fig 2A). At E11.5 *Baf60c*^-/-^ C57Bl/6 embryonic hearts had a more severe and penetrant phenotype, with a thin compact layer and fewer or less well developed trabeculae (Fig 2B), impaired atrioventricular cushion formation, and reduced atrial septum growth. In the few surviving E14.5 *Baf60c*^-/-^ mixed background embryos, ventricular free walls were much thinner than WT (Fig 2C) and the interventricular septum was disorganized, leading to ventricular septal defects. Based on the intrinsic cardiac phenotypes, we conjectured that circulatory failure and hemorrhage were the result of impaired cardiac function of *Baf60c*^-/-^ embryos.

**Fig 2.**
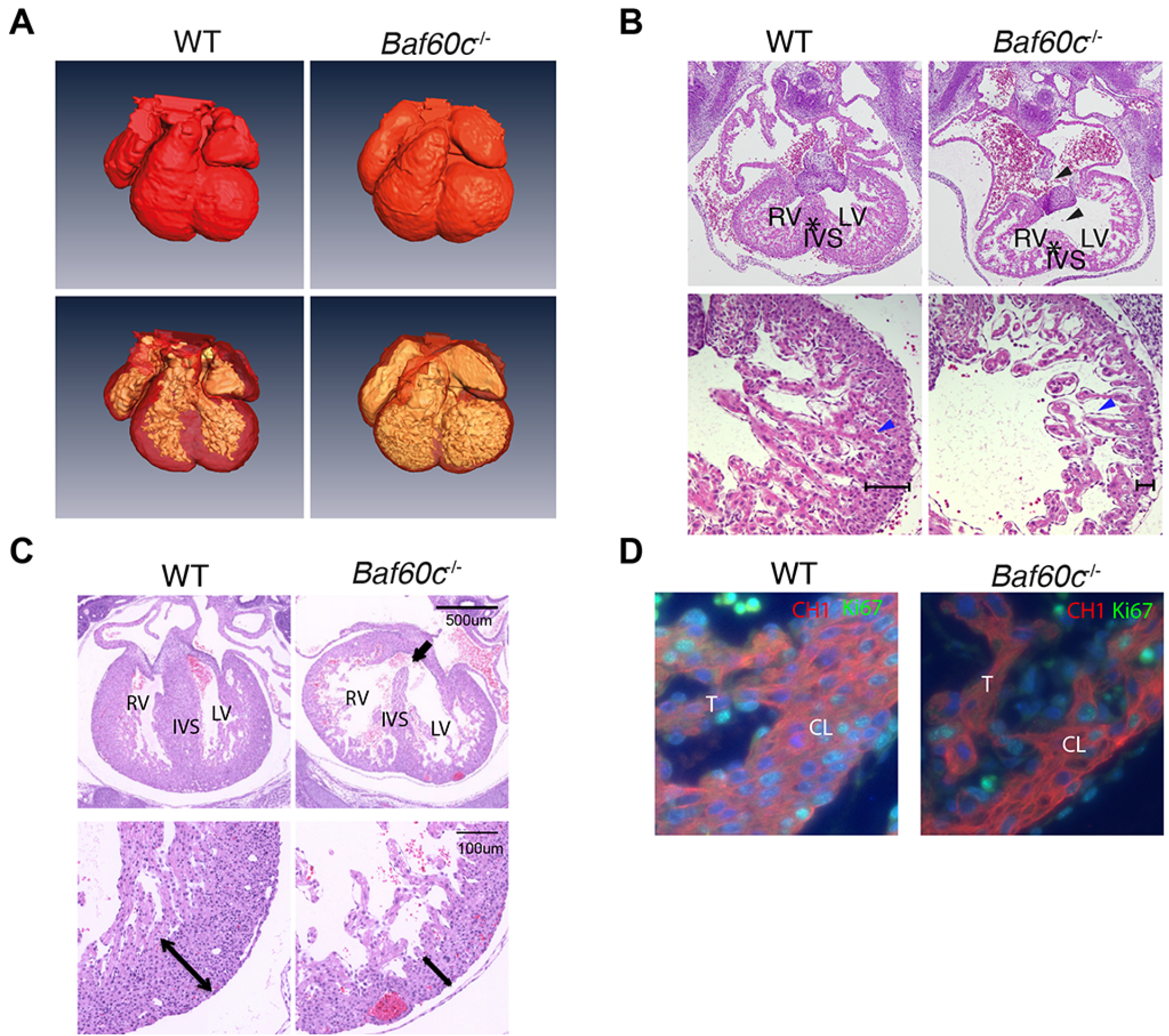
*Baf60c* deletion results in a hypoplastic embryonic heart. A: At E12.5, *Baf60c*^-/-^ embryonic hearts have similar outer dimensions as the WT, but the ventricle chambers are expanded and ventricle walls are thinner as observed by rendered OPT images. B&C: Transverse sections and HE staining of E11.5 (B) and few surviving mixed background E14.5 (C) embryonic hearts. The *Baf60c*^-/-^ hearts show incomplete inter ventricular septum formation (star), have VSDs (black arrow or arrowhead), thinner ventricle walls (brackets) and disorganized and reduced trabeculation (blue arrowhead) compared to WT strains. D: Ki67staining detects fewer proliferating cardiomyocytes in E12.5 *Baf60c*^-/-^ heart than in WT. Red: CH1 anti tropomyosin; green: Ki67. CL: compact layer. T: trabeculae.

To identify the possible cause of cardiac hypoplasia in *Baf60c* knockouts, proliferation of cardiomyocytes was assessed by staining with Ki67 antibody. Immunostaining detected fewer Ki67+ cardiomyocytes in E12.5 *Baf60c*^-/-^ ventricles than in WT (Fig 2D). Quantitation confirmed that in WT hearts, there were 32±9% Ki67+ ventricular cardiomyocytes, while in *Baf60c*^-/-^ hearts 25±5% were positive (n=4; *P*<0.05).

The embryonic heart begins to pump blood from the linear heart tube stage onwards, and its contractile function is essential for fetal life. To determine if cardiac function was affected by *Baf60c* deletion, we used high frequency ultrasound echocardiography (Zhou et al., 2002) to evaluate contractile parameters of E13.5 mixed background embryos in utero (Table 1). No regurgitation between atria and ventricles were observed in *Baf60c*^-/*-*^embryos, suggesting that cardiac valves had formed and were fully functional. However, the left ventricle fraction shortening (LVFS) of *Baf60c*^-/-^ hearts was reduced, suggesting impaired systolic function. The inter-ventricular septal fractional thickening (IVSFT) was lower than in the hearts of WT and *Baf60c*^+/-^ embryos indicating reduced myocardial contraction. The E/A ratios of *Baf60c*^-/-^ hearts for both the left and right ventricles were also significantly increased. This may indicate impaired cardiac relaxation (Zhou et al., 2003). Overall, echocardiography showed that loss of *Baf60c* affected the morphology and dimensions of the heart, and concomitantly its contractile function.

**Table 1.** High frequency echocardiography evaluation of E13.5 embryos.

|  | **WT (n=4)** | ***Baf60c*<sup>+/-</sup>**<br>**(n=5)** | ***Baf60c*<sup>-/-</sup>**<br>**(n=5)** | **P-value** |
| --- | --- | --- | --- | --- |
| **LV ESD (mm)** | 0.67±0.03 | 0.72±0.02 | 0.79±0.02 | *P*=0.0087 |
| **LV EDD (mm)** | 0.94±0.04 | 0.99±0.03 | 1.03±0.03 | NS |
| **LV FS (%)** | 28±2 | 27±1 | 23±0.02 | *P*=0.02 (WT vs KO)<br>*P*=0.01 (Het vs KO) |
| **RV ESD (mm)** | 0.64±0.03 | 0.6±0.04 | 0.69±0.03 | NS |
| **RV EDD (mm)** | 0.85±0.02 | 0.89±0.03 | 0.92±0.02 | NS |
| **RV FS (%)** | 25±3 | 32±2 | 24±3 | NS |
| **IVSTes (mm)** | 0.32±0.02 | 0.36±0.03 | 0.25±0.03 | *P*=0.0023 |
| **IVSTed (mm)** | 0.22±0.03 | 0.25±0.03 | 0.20±0.02 | NS |
| **IVSFT** | 40±4 | 41±4 | 26±6 | *P*=0.004 (WT vs KO)<br>*P*=0.006 (Het vs KO) |
| **Mitral peak E/A** | 0.12±0.04 | 0.25±0.05 | 0.32±0.04 | *P*=0.02 |
| **Tricuspid peak**<br>**E/A** | 0.19±0.03 | 0.17±0.03 | 0.34±0.04 | *P*=0.008 |

IVSFT: inter-ventricular septal fractional thickening; IVSTed: end-diastolic inter ventricular septum thickness; IVSTes: end-systolic inter-ventricular septum thickness; IVFST:; LV EDD: left ventricular end-diastolic diameter; ESD: LV ESD: left ventricular end-systolic diameter; LV FS: left ventricular fractional shortening; RV EDD: right ventricular end-diastolic diameter; ESD: RV ESD: right ventricular end-systolic diameter; RV FS: right ventricular fractional shortening. Peak E/A: the ratio of peak velocities of the early diastolic waveform (E wave) to the late diastolic waveform during atrial contraction (A wave) at either mitral or tricuspid orifices.

We assessed the tissue-specificity of the *Baf60c*^-/-^ phenotype by crossing *Baf60c*^*flox*/*flox*^ mice with *Nkx2-5::Cre* mice. *Nkx2-5::Cre* deletes loxP-flanked DNA from E8.0 in all cardiac precursors (Moses et al., 2001). *Nkx2-5::Cre*::*Baf60c*^*flox*/-^ mice had morphological defects similar to those found in the least severely affected *Baf60c*^-/-^ embryos (Fig 3), indicating that the constitutive null phenotype reflects primary loss of *Baf60c* in the developing heart, but also potentially an earlier function in precursors that do not yet express *Nkx2-5* (Devine et al., 2014).

**Fig 3.**
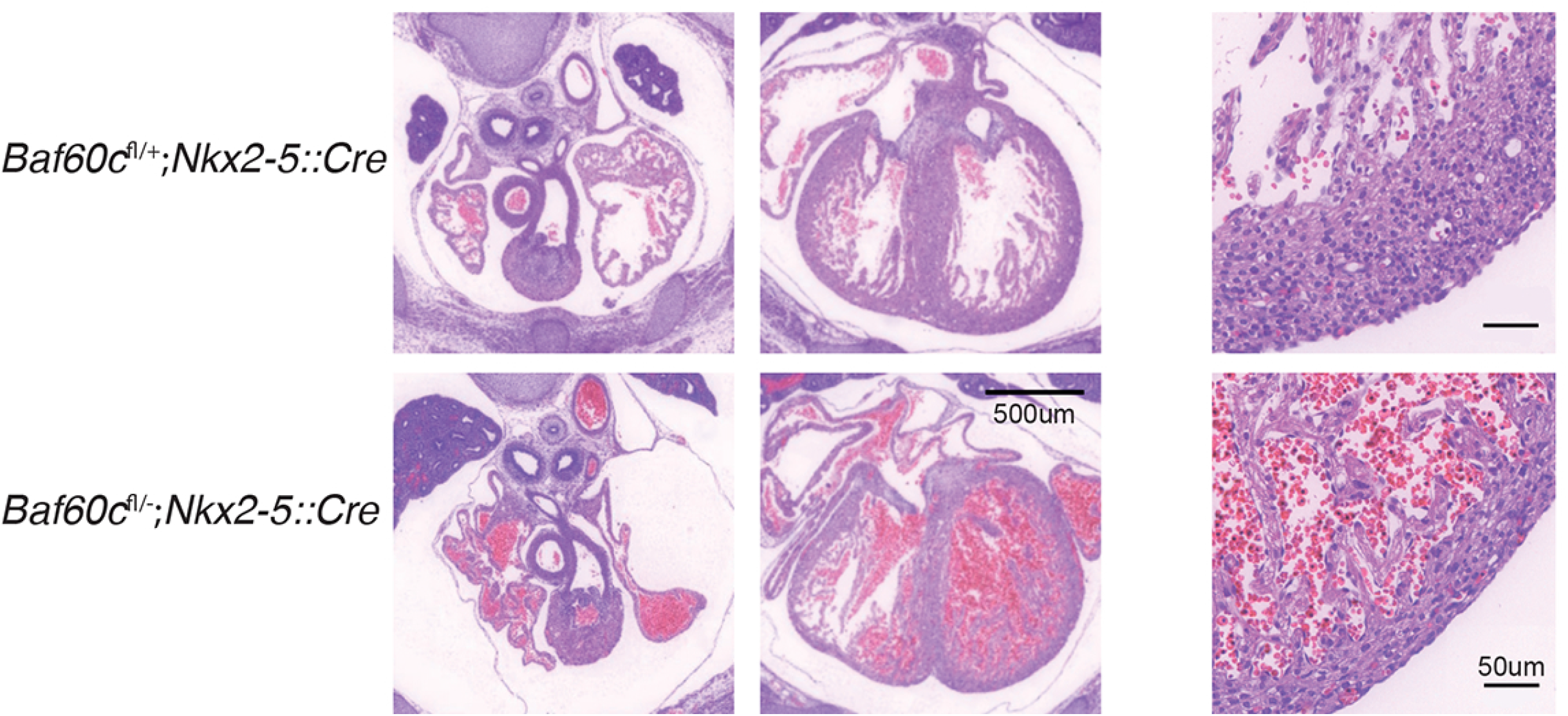
Deletion of Baf60c with *Nkx2-5*^*Cre*^. HE staining of transverse sections of E14.5 mouse heart shows thinner myocardium, reduced trabeculation, and ventricular septation defects.

### Loss of Baf60c in cardiomyocytes results in postnatal cardiomyopathy

After birth, heart development switches from cell proliferation to hypertrophic growth. The structure and physiological function of the myocardium undergo a series of changes to adapt to a new hemodynamic environment. We deleted *Baf60c* in the myocardium at later developmental stages by crossing the *Baf60c*^*flox*/*flox*^ allele with *Myh6::Cre* (Agah et al., 1997). This manipulation bypassed the embryonic lethality of the constitutive deletion, as *Baf60c*^*flox*/-^;*Myh6::Cre* (*Baf60c*^*Myh6KO*^) mice were born alive and showed no obvious morphological changes before postnatal day (P) 7. After P7, some of the *Baf60c*^*Myh6KO*^ pups were growth-delayed compared with their littermates and died before weaning. Other *Baf60c*^*Myh6KO*^ mice survived after weaning without obvious morphological defects, but at ~4 to 6 weeks exhibited symptoms of heart failure, including weight loss, reduced activity level, hunched back and labored breath. The remaining *Baf60c*^*Myh6KO*^ mice appeared normal, but died suddenly. All *Baf60c*^*Myh6KO*^ mice died before 4 months of age (Fig 4A).

**Fig 4.**
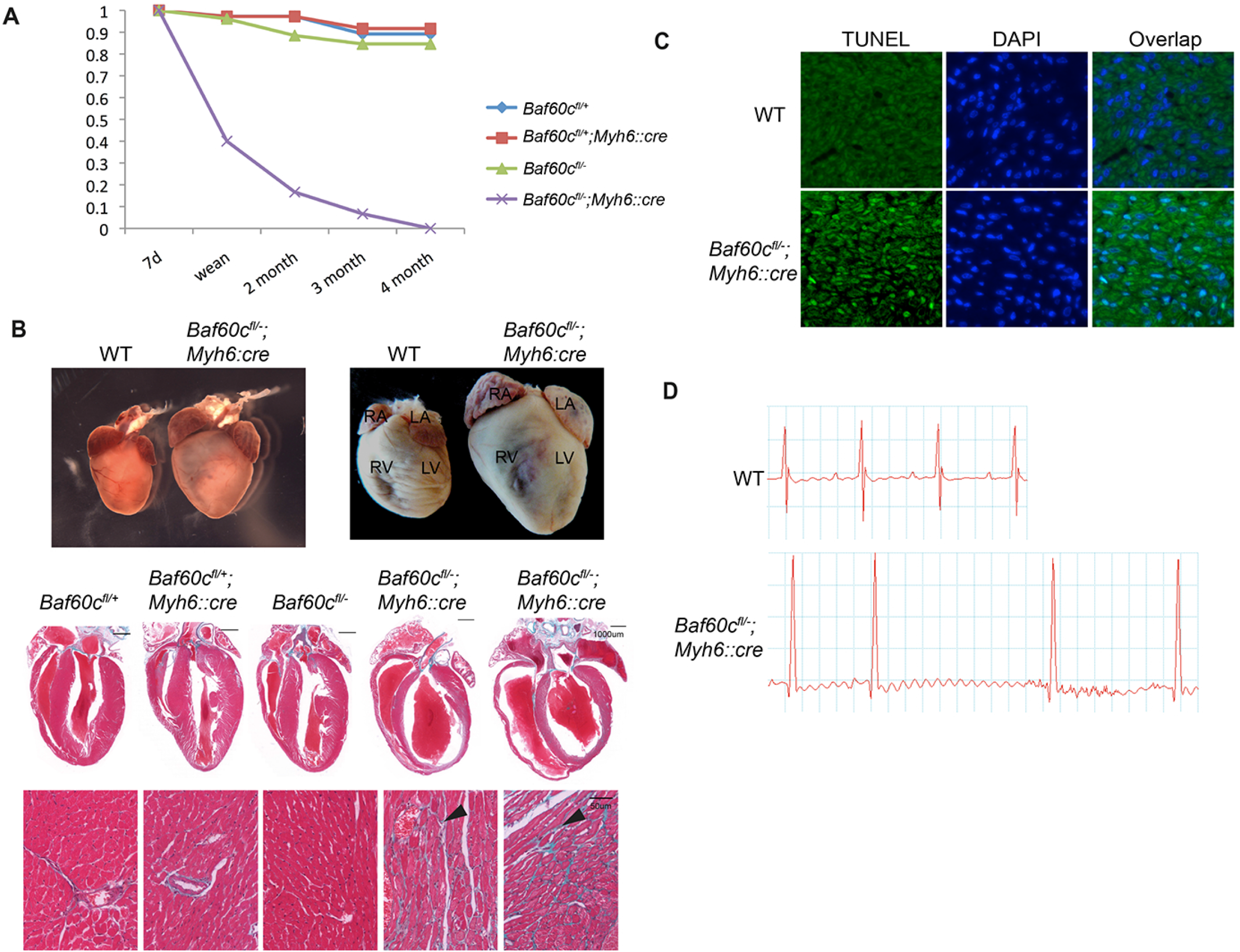
Deletion of *Baf60c* in myocardium results in dilated chambers and impaired cardiac function. A: *Baf60c*^*fl/-*^*;Myh6::Cre* mice all die before 4 months of age. B: *Baf60c*^*fl/-*^*;Myh6::Cre* mice have enlarged hearts and dilated chambers, as shown with whole-mount (top panel) and 4-chamber view sections (middle panel). Left panel: P10. Right panel: 8 week hearts. Masson Trichrome staining detects fibrosis in ventricle myocardium (bottom panels, arrowheads). C: *Baf60c*^*Myh6KO*^ myocardium have high level apoptosis. Green: TUNEL. Blue: DAPI. D: Representative electrocardiogram of adult WT and *Baf60c*^*fl/-*^*;Myh6::Cre* mice.

To investigate the reason for the early mortality in *Baf60c*^*Myh6KO*^ mice, their hearts were dissected at different ages for morphology and histology analysis. At all the observed stages (P10, P21 and 8 weeks), the hearts of *Baf60c*^*Myh6KO*^ mice were enlarged compared with the controls (Fig 4B). Histology revealed chamber dilation (Fig 4B). Masson’s trichrome staining detected broad myocardium interstitial fibrosis in the *Baf60c*^*Myh6KO*^ myocardium, while this was not observed in any other genotypes (Fig 4B, lower panels. A high level of apoptosis was also detected in myocardium of adult *Baf60c*^*Myh6KO*^ mice (Fig 4C).

The chamber dilation and fibrosis observed in the hearts of *Baf60c*^*Myh6KO*^ mice raised the question of whether cardiac function was also affected. We measured cardiac contractile function of 8-week old mice that lacked outward signs of heart failure or growth delay, using high frequency echocardiography (Table 2, n=6). Confirming the histological results, the left ventricles of *Baf60c*^*Myh6KO*^ mice were prominently dilated, and the anterior and posterior ventricle walls of *Baf60c*^*Myh6KO*^ mice were thinner and the chamber contraction ratio decreased. The aortic time-velocity integral (TVI, which measures the distance traveled by a volume of blood during a time interval) increased, probably because of the enlarged ventricle volume. The fraction shortening (FS) and cardiac output was reduced, consistent with the cardiac failure symptoms of *Baf60c*^*Myh6KO*^ mice. We performed electrocardiogram analysis to measure the conduction function of *Baf60c*^*Myh6KO*^ mice (Fig 4D, Table 3, n=5-6). Compared with other genotypes, *Baf60c*^*Myh6KO*^ mice had significantly slower heart rates, shortened conduction time through the AV node (PR interval), and prolonged QRS duration, suggesting longer depolarization-repolarization time of the ventricle. P height, which indicates atrial depolarization, was reduced. Thus, clear and significant conduction defects accompany contractile deficiency in *Baf60c*^*Myh6KO*^ mice.

**Table 2.**
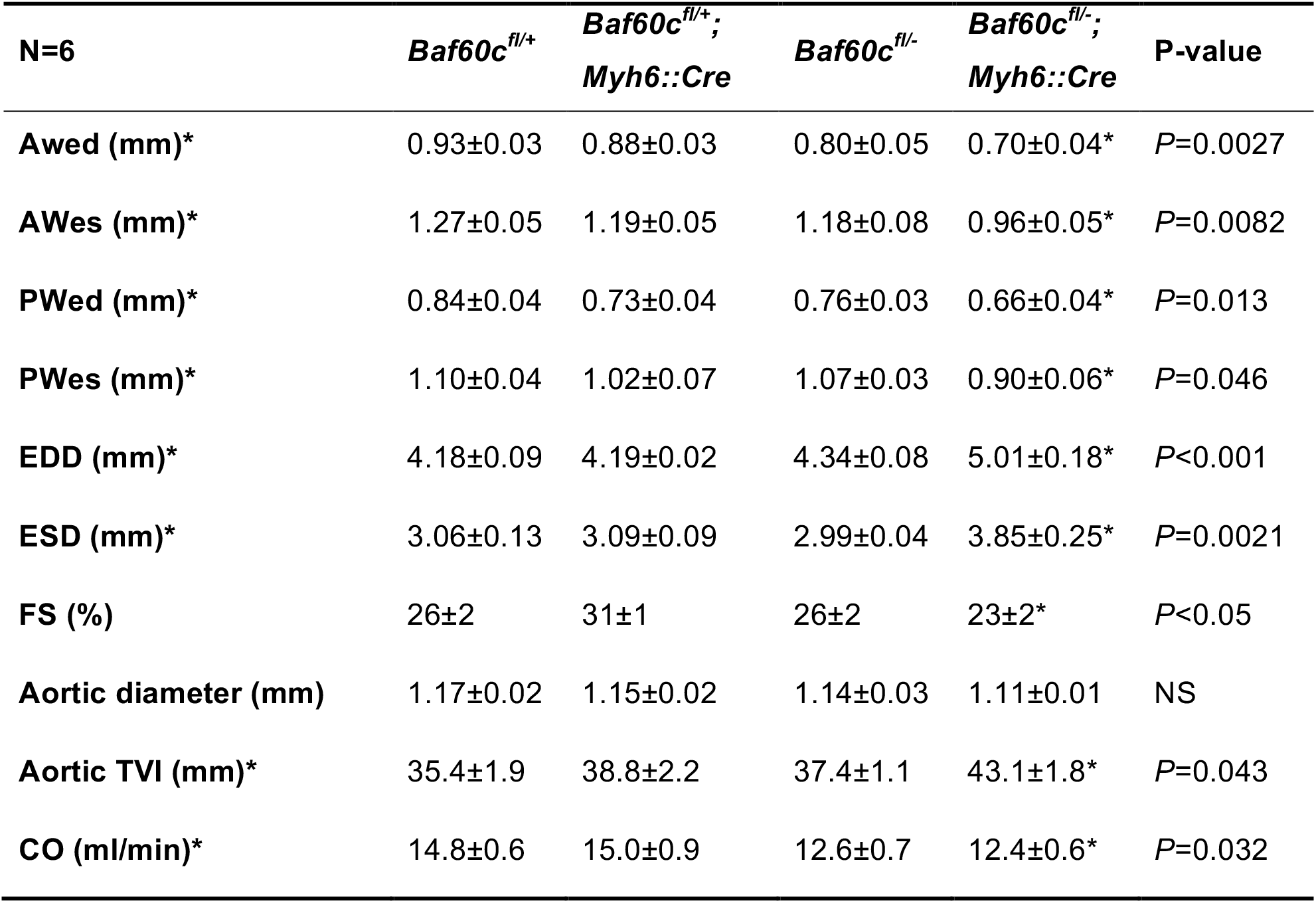
Echocardiography analysis of the cardiac contractile function *Baf60c*^*Myh6KO*^ mice at 8 weeks of age.

AWed: anterior wall thickness at end diastole; AWes: anterior wall thickness at end-systole; EDD: end-diastolic diameter; ESD: end-systolic diameter; FS: fractional shortening; PWed: posterior wall thickness at end-diastole; PWes: posterior wall thickness at end systole; TVI: time-velocity integral; CO: cardiac output. Asterisks indicate significantly different values.

**Table 3.** Electrocardiogram analysis of *Baf60c*^*Myh6KO*^ mice.

| **N=6** | ***Baf60c*<sup>fl/+</sup>** | ***Baf60c*<sup>fl/+</sup>;**<br>***Myh6::Cre*** | ***Baf60c*<sup>fl/-</sup>** | ***Baf60c*<sup>fl/-</sup>;**<br>***Myh6::Cre*** | **P value** |
| --- | --- | --- | --- | --- | --- |
| **Heart rate** | 558±17 | 560±14 | 521±11 | 453±18* | *P*=0.036 |
| **P duration (ms)** | 10.2±1.0 | 7.7±0.7 | 9.8±0.6 | 9.2±0.7 | NS |
| **PR duration (ms)** | 36.5±0.8 | 36.5±0.5 | 34.7±0.4 | 30.6±0.5* | *P*=0.019 |
| **QRS duration (ms)** | 7.8±0.2 | 8.0±0.3 | 8.2±0.2 | 11.0±0.5* | *P*<0.0001 |
| **QT duration (ms)** | 21.3±0.5 | 20.0±0.5 | 20.0±0.6 | 25.0±0.5* | *P*=0.031 |
| **QT Max Duration (ms)** | 8.7±0.7 | 8.00±0.7 | 8.17±0.2 | 11.0±0.5* | *P*=0.005 |
| **P height (mV)** | 0.15±0.02 | 0.10±0.03 | 0.13±0.03 | 0.07±0.01* | *P*=0.015 |
| **QRS height (mV)** | 1.1±0.2 | 1.0±0.2 | 1.8±0.2 | 2.2±0.1* | *P*<0.001 |

Asterisks indicate significantly different values.

### Myofibrillar defects of Baf60c KO cardiomyocytes

The cardiac structural and functional defects in *Baf60c*^-/-^ are a reflection of an underlying cellular defect. To address this, we used electron microscopy to observe cardiomyocyte ultrastructure. At E12.5, sarcomeres of *Baf60c*^-/-^ hearts were disarrayed, and the thick and thin filaments were discontinuous and poorly aligned. Z-disks were loosely packed and did not have clear defined borders as was found in WT sarcomeres. The I band (thick-filament free zone) and the M bands (myosin head free zone of the thick filaments) located in the middle of sarcomere were almost undetectable (Fig 5A, top panel). Similar defects also existed in adult *Baf60c*^*Myh6KO*^ cardiomyocytes. We measured sarcomere length in adult hearts and found the sarcomere length (the length between two adjacent Z-disks) of *Baf60c*^*Myh6KO*^ mice was significantly shorter than WT (Fig 5B).

**Fig.5.**
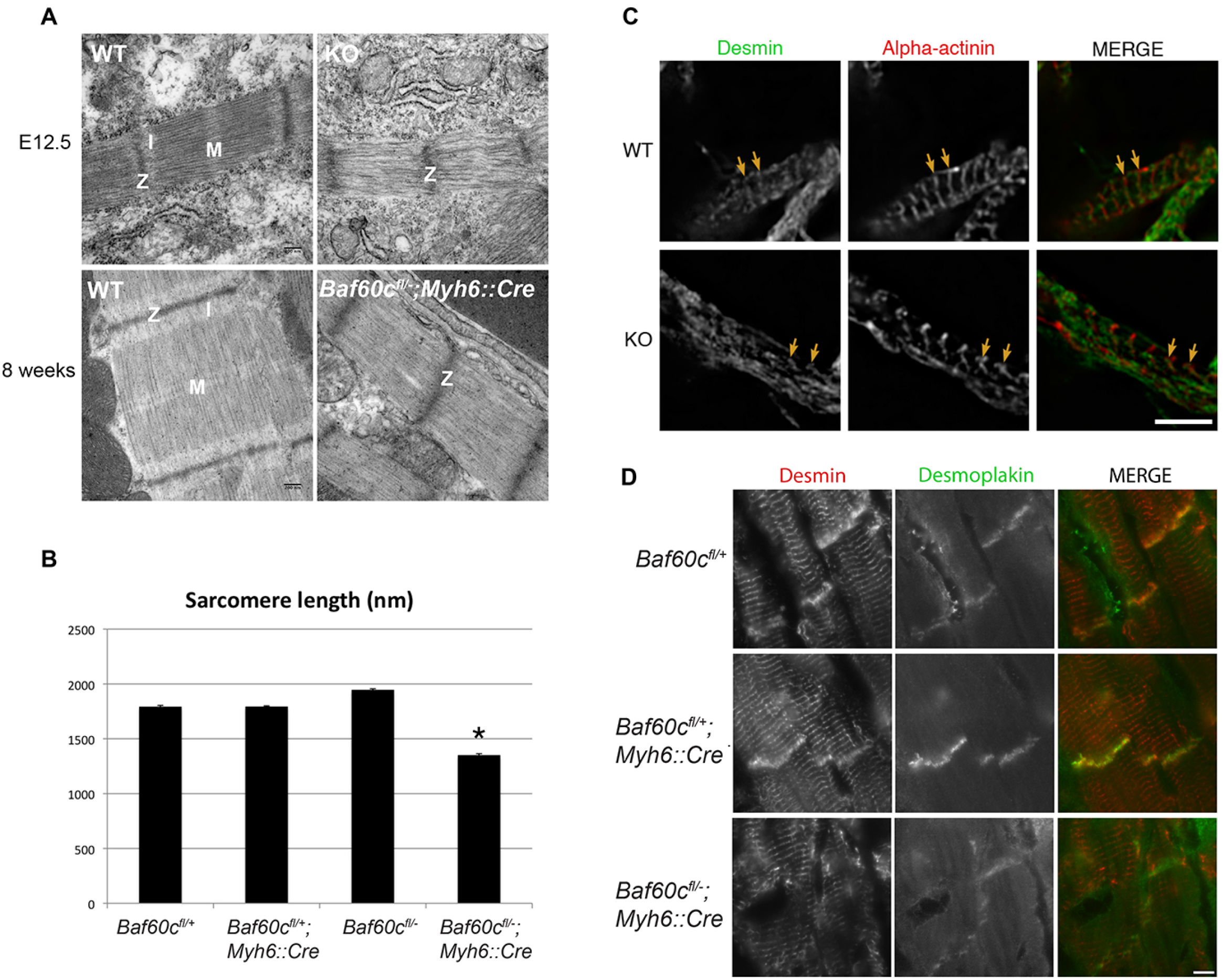
Myofibrillar defects of *Baf60c*^-/-^ cardiomyocytes. A: Cardiomyocyte ultrastructure of WT and *Baf60c*^-/-^ under Transmission electron microscopy (TEM). Z: Z-disk. I: I-band; M: M-line. B: In adult mice, sarcomeres of *Baf60c*^*fl/-*^*;Myh6::Cre* cardiomyocytes are shorter. Mouse hearts were not relaxed before sample preparation, but only relaxing sarcomere were measured. C and D: localization of Desmin was disturbed in embryonic and adult hearts in the absence of Baf60c.

We examined the distribution of several important structural proteins in cardiomyocytes by immunofluorescence deconvolution microscopy, and found that the localization of Desmin in Z-disks of embryonic cardiomyocytes was disturbed in *Baf60c*^-/-^ hearts (Fig 5C). In adult *Baf60c*^*Myh6KO*^ hearts, localization of Desmin in intercalated discs was also reduced (Fig 5D), and the pattern of Desmin localization was perturbed (poorly aligned). These observations are similar to what was observed by electron microscopy and together, showing disrupted myofibril alignment and sarcomere structure in the absence of *Baf60c*.

### Cardiac gene expression program regulated by Baf60c

To identify genes regulated by Baf60c, we used RNAseq to analyze total RNA prepared from *Baf60c*^-/-^ hearts and control hearts harvested at E10.5 and E12.5, and from P7 *Baf60c*^*Myh6KO*^ and control hearts. We identified 788 genes that were differentially expressed by at least 1.25 fold (p<0.05) in at least one stage versus wild type (Fig 6A). Among all the genes and all the analyzed stages, there were 132 genes upregulated and 175 down-regulated at all time points. Misregulation of major cardiac transcription factors or signaling molecules was not observed. Instead, consistent with the ultrastructural findings, many genes related with cardiac metabolism and striated muscle contraction such *Acta1*, *Aldh1l2*, *Casq1*, *Casq2*, *Ckm*, *Ckmt2*, *Trim72*, *Kbtbd10 (Krp1)*, *Myh7b*, *Myl3, Mylpf, Obscn,* and *Tnni2* were identified as downregulated in embryonic and adult *Baf60c*-deficient hearts (Fig 6B). A broader range of cardiac function related genes was deregulated in the *Baf60c*^*Myh6KO*^ hearts, including *Gja3*, *Myl1*, *Myl4*, *Myl7*, and *Tnni1*. The postnatal deletion of *Baf60c* also resulted in induction of *Nppa*, as might be expected in a cardiomyopathic heart (Houweling et al., 2005), but the induction was mild (only 2-fold increase), indicating a potential deficiency in upregulation of this marker of cardiac stress. In fact, the usual set of cardiac stress-responsive genes was not present in the *Baf60c*^*Myh6KO*^ cardiac gene expression program. Gene ontology (GO) analysis of genes repressed by *Baf60c* in postnatal heart enriched for biological processes involved in broad developmental processes and extracellular structure organization (Fig 6C). On the other hand, *Baf60c* activated genes were enriched for muscle system processes, regulation of muscle cell differentiation, muscle contraction, and sarcomere and actin cytoskeleton organization (Fig 6D). An enrichment of cell cycle-related genes was also apparent; it is not clear what this signifies, and may reflect a role for *Baf60c* in regulating perinatal proliferation, which was not addressed in this study. These results collectively suggest that *Baf60c* is required for proper expression of genes encoding components or regulators of the contractile apparatus.

**Fig 6:**
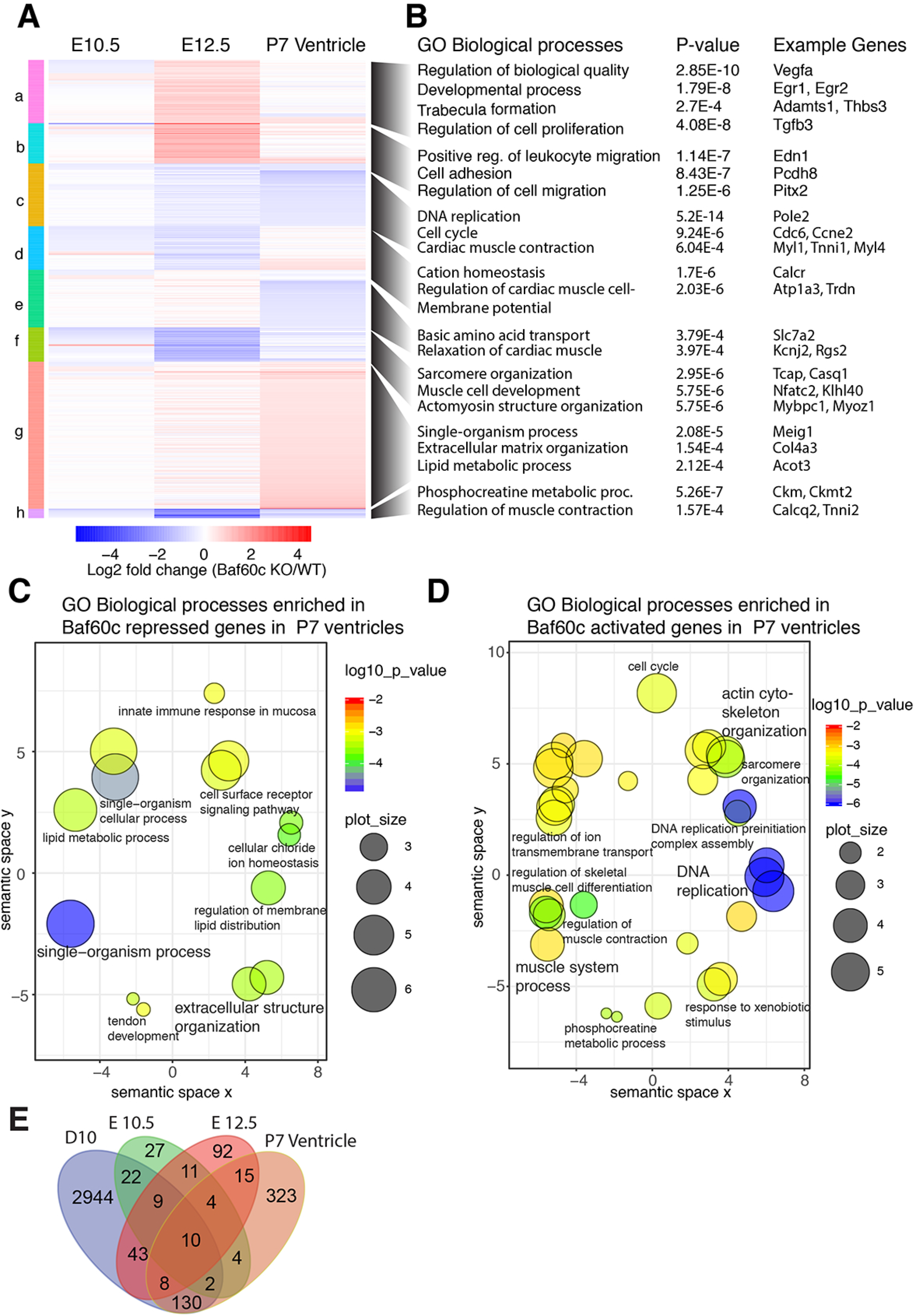
BAF60c transcriptionally affects cardiac morphogenesis and function. A: Heat map comparing genes affected by *Baf60c* loss in embryonic hearts (E10.5 and E12.5) and postnatal ventricles (postnatal Day (P) 7). Significantly affected (≥1.25 fold, p<0.05) genes in at least any one stage were selected and clustered. B: Gene ontology (GO) biological processes enriched in each of these clusters and example genes in that category are shown. C&D: Genes repressed (C) or activated (D) by Baf60c in P7 ventricles were analyzed for enrichment of GO biological processes and are plotted. The color of circles represents p-value of enrichment and size represent the size of the GO term. E: Venn diagram showing genes mis-regulated in absence of Baf60c in embryonic heart at E10.5, E12.5 and P7 Ventricle and compared cardiac myocyte (Day 10) stages of in-vitro directed cardiac differentiation.

The analysis of gene expression in whole hearts has the disadvantage that a heterogeneous mix of cells may prevent the clear identification of the full set of Baf60c-regulated genes, and also that some changes in gene expression may be secondary to altered hemodynamics. We compared the set of genes altered in the Baf60c mutant hearts with RNAseq analysis of cardiac precursors and cardiomyocytes differentiated in vitro from WT and *Baf60c*^-/-^ embryonic stem (ES) cells (Hota et al., 2017). Considerable overlap was found for the E12.5 KO hearts and ES cell-derived cardiac precursors differentially expressed genes, and more significant overlap was found for E12.5 KO and *Baf60c*^*Myh6KO*^ hearts with ES cell-derived cardiomyocyte (Fig.6E). These comparisons show that both in vitro and in vivo, *Baf60c* regulates a set of genes important for cardiac morphogenesis and function.

### Baf60c functionally interacts with Myocardin

We previously identified TBX5 and NKX2-5 as potential BAF60c-interacting proteins (Lickert et al., 2004). Here we demonstrate by GST pulldown that these interactions can be direct (Fig 7A). We mapped the BAF60c interaction domain to an N-terminal region (aa residues xx-xx) that contains a nuclear localization signal sequence (Fig 7B). To further elucidate the molecular mechanism of BAF60c function, we searched for potential association partners of BAF60c. In a yeast 2 hybrid screen of a human heart cDNA library, using BAF60c as the bait, we identified few potential interacting factors (BAF155, FEZ1, MYOCD). BAF155 is a component of the BAF complex, which indicates a direct interaction between these two BAF complex subunits. Of particular interest amongst candidate interactors was Myocardin (MYOCD), a transcriptional co-factor of SRF and MEF2c (Creemers et al., 2006; Wang et al., 2001). GST pull-down assay between GST-fused BAF60c and in-vitro synthesized MYOCD confirmed the direct association, and mapped the association domain of MYOCD with BAF60c to amino acids 328-554 (Fig 7C). *Myl1* is a bona fide direct target of MEF2c/Myocardin (Creemers et al., 2006) and was downregulated in the absence of *Baf60c*. In an in vitro promoter activation assay, BAF60c could potently enhance the activation of the *Myl1* promoter by MYOCD and MEF2c (Fig 7D). Together our data suggest that BAF60c can function as a partner of MYOCD in cardiac development, and that this interaction may be important for the activation of a gene expression program essential for the fundamental functional properties of cardiomyocytes.

**Fig 7:**
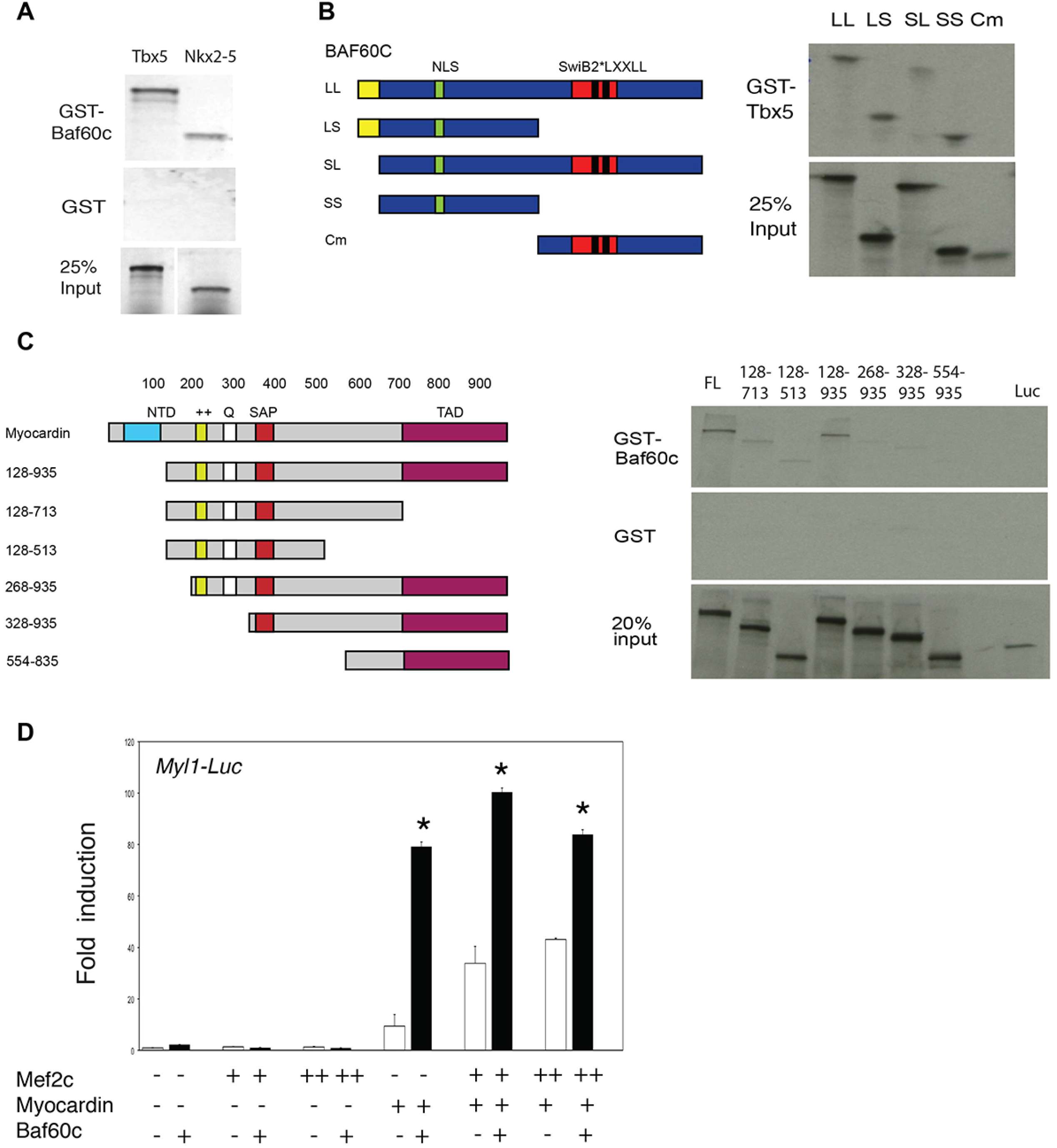
Interaction between BAF60c and cardiac transcription factors. A. GST-fused Baf60c associates with ^35^S labeled, in-vitro synthesized TBX5 and NKX2-5. B. BAF60c associates with TBX5 through its N-terminal domain. Left: schemes representing serial deletion constructs of BAF60c. Right: Mapping the BAF60c associating domain with GST-fused TBX5. C. BAF60c associates with full length Myocardin and serial deletions. Left: Schemetic representation of myocardin deletion constructs. The region between Myocardin 328-554 aa was essential for association with Baf60c. D. BAF60c enhances activation of MYOCD and MEF2c on the *Myl1* luciferase reporter.

## Discussion

We have demonstrated the requirement for *Baf60c* in cardiomyocyte function throughout heart development. Loss of *Baf60c* both prenatally and postnatally resulted in cardiac hypoplasia and defective heart function. *Baf60c* regulates programs of gene expression that are essential for primary functions of cardiomyocytes, including broad sets of genes essential for sarcomere function and cardiac metabolism.

The *Baf60c* constitutive knockout phenotype is milder than the mouse shRNA knockdown phenotypes published previously (Lickert et al., 2004). The shRNA knockdowns were performed using two different shRNAs, minimizing the possibility of off-target effects, and the phenotype was rescued by over-expression of BAF60b, indicating significant specificity of the shRNAs. A similar discrepancy exists for *Ifitm* genes, for which the shRNA phenotype is more severe than that of a genetic deletion (Lange et al., 2008; Tanaka et al., 2005). The possible reasons for the different phenotypes between the shRNA and the genetic null might include effects compounding the loss of *Baf60c* function such as overloading of the microRNA processing machinery by overexpressing shRNAs at high levels, other non-specific effects inherent to overexpression of shRNAs in the mouse embryo, or failure to compensate for immediate repression of gene function by RNAi. The genetic deletion presented here confirms an important role for *Baf60c* in heart development, and extends these findings significantly.

The phenotype resulting from loss of *Baf60c* suggests that *Baf60c* has a specific role in regulating gene expression programs necessary for cardiac growth and contractile function. BRG1, the core ATPase of BAF complexes, has broad and critical roles in supporting cardiomyocyte proliferation and differentiation at embryonic stages and hypertrophic growth in the stressed adult heart (Hang et al., 2010; Takeuchi et al., 2011). Our data suggest that a BAF60c-containing cardiac-specific BAF complex has a more specialized role, and may have evolved to provide fine-tuned and specific gene regulation in the mammalian heart. Indeed, we have isolated BAF complexes during in vitro cardiac differentiation and have identified that BAF60c-containing complexes in cardiomyocytes have a composition that differs from many BRG1-containing complexes (Hota et al., 2017). In skeletal muscle differentiation, BAF60c interacts with MYOD to activate muscle-specific genes (Forcales et al., 2012), and is essential for HDAC-dependent fibro-adipogenic precursor differentiation in dystrophic muscle (Saccone et al., 2014). The set of genes that are altered due to depletion of BAF60c in differentiating C2C12 cells (Forcales et al., 2012) is remarkably similar to those altered by loss of BAF60c in the heart, indicating a commonality in the regulatory program controlled by BAF60c in cardiac and skeletal muscle. The role of BAF60c in glycolytic metabolism of fast-twitching muscle has also been described (Meng et al., 2013); whether Baf60c has a specific function regulating metabolic switching during cardiomyocyte maturation will be a potential direction for future studies.

*Myocd* is an essential factor for embryonic cardiac gene expression and postnatal myocardial function (Creemers et al., 2006) (Huang et al., 2012; Huang et al., 2009). Loss of *Myocd* in cardiac precursors results in a phenotype very similar to that of *Baf60c*-null embryos, with death around E13.5, thinned myocardium, ventricular septal defects, and reduced proliferation (Huang et al., 2012). Cardiomyocyte-specific deletion of *Myocd*, as with that of *Baf60c*, also results in sarcomere disorganization, mislocalization of Desmin, and apoptosis (Huang et al., 2009). However, the changes in gene expression documented in *Myocd* deficient hearts are not fully recapitulated by the loss of *Baf60c*, indicating that a MYOCD/BAF60c interaction may target a specific subset of *Myocd*-regulated genes, such as *Myl1* and others. The association domain of MYOCD with BAF60c did not differentiate between the smooth muscle and cardiac isoforms (Creemers et al., 2006), suggesting BAF60c can either associate with both and regulate different programs, or there are other mechanisms *in vivo* controlling selective association with either isoform. BAF60c can act on SRF-dependent promoters to regulate smooth muscle gene expression (Sohni et al., 2012); it remains to be determined whether this activity also involves an interaction with Myocardin.

Mutations in many cardiac transcription factor and structural genes have been identified to result in congenital heart defects and cardiomyopathy (Ahmad et al., 2005; Bruneau, 2008; Fahed et al., 2013). Mutations in histone modifying complex subunit genes and in some chromatin remodeling protein-encoding genes have been identified in patients with CHDs (Homsy et al., 2015; Zaidi et al., 2013). While no mutations in *SMARCD3*, which encodes BAF60c, have been associated with CHDs, the functional interaction of BAF60c with several transcription factors implicated in CHDs suggests that a potential underlying mechanism for CHDs may be dependent on BAF60c. Indeed, our recent proteomic analysis of BAF complexes has identified WDR5, mutated in human CHD, as part of a cardiac-enriched BAF complex (Hota et al., 2017). In conclusion, we have demonstrated the essential role of BAF60c in cardiac growth and function, and implied a possibility of chromatin remodeling factors contributing to CHDs.

## Methods

### ES cell targeting and mouse line establishment

A *Baf60c* genomic DNA fragment with loxP sites flanking 1^st^ exon to 4^th^ exon and Frt-Neo-Frt cassettes downstream of 4^th^ exon was constructed using bacterial recombineering (Fig. 1A). For gene targeting, 5x10^6^ R1 ES cells were trypsinized and electroporated with 25ug linearized targeting DNA. The electroporated cells were selected with 160ug/ml G418 (Gibco # 10131) for 7 days. Correctly targeted clones were identified using Southern Blot with DNA probes located outside the targeting DNA and labeled with ^32^P (Perkin Elmer). The clones were then expanded and used for diploid aggregation. High ESC contributed chimera males were bred with ICR and C57/BL6 for germline transmission. *Baf60C*^*neo*/+^ progeny were mated with FLPe-expressing mice (B6;SJL-Tg(ACTFLPe)9205Dym/J, maintained at the Toronto Center for Phenogenomics, TCP) to remove the *Neo* cassette between the *frt* sites and yield *Baf60c*^*flox*/+^ mice. To generate the *Baf60c* deletion, *Baf60c*^*flox*/+^ mice were mated with pCX-NLS-Cre mice (maintained at the TCP).

### Mouse and embryo genotyping

The *Baf60c*^*flox*/+^ and *Baf60c*^+/-^ mice were genotyped by PCR using 3 primers: WTfor (5’-CGTTCTGCAAGATGGTCTGA-3’), DELfor (5’-AGGCAGACCCAAGCTTGATA-3’) and Rev (5’-CATCAGAGTCTTCCGCATCA-3’). Baf60c deletion band is 250bp, wild type is 350bp and Baf60cfloxed is 470bp. Postnatal mouse tissues (tail tips or ear notches) and embryo tissues (yolk sac, tails, limb buds) were prepared with the tissue preparation buffer of the Sigma Extract-N-Amp tissue PCR kit (Sigma, XNAT2).

### Histology

Mouse embryos or tissues were fixed with 4% PFA, dehydrated and embedded with paraffin and sectioned into 4μm sections then mounted on glass slides. The slides were then stained using standard histology protocols.

### Whole mount In-situ hybridization

Whole mount in-situ hybridization on mouse embryos from E7.5 to E10.5 was performed according to standard protocols using the *Baf60* in situ hybridization probe (Lickert et al., 2004).

### Optical projection tomography

Optical projection tomography (OPT) was performed as described previously (Sharpe et al., 2002) using a OPT system built in-house. E12.5 embryos were harvested, genotyped, fixed with 4% PFA overnight and washed with PBS. The specimens were then embedded in 1% low melting point (LMP) agarose and subsequently cleared using a 1:2 mixture of benzyl alcohol and benzyl benzoate (BABB). The index-matched specimen was suspended from a stepper motor and immersed in a BABB bath encompassed in a glass cuvette. Light from a mercury lamp was directed onto the specimen and filter sets were used to create fluorescent images of the specimen. An autofluorescence projection was captured with using a GFP filter set in the illumination and detection light path. Images of the specimen were formed using a Qioptiq Telecentric Zoom 100 microscope equipped with a 0.5X OPTEM objective lens. Projection images were acquired with a Retiga-4000DC CCD camera with pixel size equal to 7.4um/pixel. The sample was rotated in finite steps, 0.3 degrees, through a complete revolution totaling 1200 projections. Image reconstruction into a 3D data set was then executed by a modified Feldkamp algorithm in supplied software by SkyScan (Nrecon). The resultant OPT images have an isotropic 8.8 micron pixel size.

### RNA-seq

Mouse embryos from *Baf60c*^+/-^ intercross timed pregnancy at E10.5 and E12.5, or ventricles from *Myh6::Cre;Baf60c*^*fl*/+^ X *Baf60c*^+/-^ intercrosses were harvested. Their hearts were individually dissected and snap-frozen with liquid nitrogen. RNA was prepared from each single heart using PicoPure RNA Isolation kit (Arcturus). RNA quantity and quality was analyzed using Agilent RNA 6000 Nano Kit. RNA-seq was performed as described (Christodoulou et al., 2011; Christodoulou et al., 2014). RNA reads were aligned with TopHat / Bowtie and Useq was used for the analysis of differential expression. RNAs that showed significant differential expression between wild type and *Baf60c*^-/-^ (p-value <0.05) and also changed more than 1.25 fold in *Baf60c*^-/-^ over wild type at a specific stage of differentiation were selected for analysis, avoiding duplicate and redundant entries.

### Transmission Electron Microscopy

Mouse E10.5, E12.5 embryonic hearts and 8 week old adult hearts were dissected. For embryonic hearts, the whole heart was used for fixation and section. For adult hearts, pieces of 3~4 mm in size cut from the left ventricle were used as specimens. Pieces of specimen were fixed in a fixative containing 4% formaldehyde and 1% glutaraldehyde in phosphate buffer, pH7.3, and then post fixed in 1% osmium tetroxide. The specimens were then dehydrated in a graded series of acetone from 50% to 100% and subsequently infiltrated and embedded in Epon-Araldite epoxy resin. The processing steps from post fixation to polymerization of resin blocks were carried out in a microwave oven, Pelco BioWave 34770 (Pelco International, CA) using similar procedures, with slight modification, as recommended by the manufacturer. Ultrathin sections were cut with a diamond knife on the Reichert Ultracut E (Leica Inc., Austria). Sections were stained with uranyl acetate and lead citrate before being examined in the JEM-1011 (JEOL USA Corp., Peabody, MA). Digital electron micrographs were acquired directly with a 1024 X1024 pixels CCD camera system (AMT Corp., Denver, MA, USA) attached to the TEM.

### Echocardiography assessment of cardiac functions

E13.5 embryos were analyzed with a Vevo770 ultrasound machine (VisualSonics). Pregnant *Baf60c*^*+/-*^ female mice carrying the embryos at the required developmental stages were examined under isoflurane anaesthesia. The uterus were exposed from the incision and scanned with a 30 MHz transducer as described (Lickert et al., 2004). To minimize potential impairment of embryonic physiology, only 2 or 3 embryos were scanned for each female, taking about 1 hour. The mother’s heart rate was monitored throughout the scanning. For each embryo, the blood flow speed near the mitral and tricuspid valves and aorta was recorded at B-mode. The depth of ventricle walls and ventricle septation was measured at M-mode. After scanning, the embryos were harvested and genotyped. 4-5 embryos of each genotype were measured. Adult mice were analyzed using a Vevo2100 ultrasound machine (VisualSonics). The 7-8 week old animals were anaesthetized and scanned with a 30 MHz transducer as described (Zhou et al., 2005). E and A peaks in the left ventricle were measured at B-mode. The chamber dimensions and ventricle wall depths as well as ventricle septation depth were measured at M-mode. For each genotype, 5-6 mice were measured.

### Electrocardiography

Mice were anesthetized with 1-2% isoflurane and lead II ECG was recorded from needle electrodes inserted subcutaneously into the right forelimb and into each hindlimb. The signal was recorded for ~1 minute. The ECG was recorded with Power Lab/4SP (AD Instruments) and analyzed using the SAECG (signal-averaged electrocardiogram) extension for Chart 4 (v4.2.3 for Macintosh, AD Instruments).

### Immunofluorescence Microscopy

Sarcomeric architecture and organization were assessed in E12.5 and adult hearts via double immunofluorescence staining. Heart tissue was embedded in Tissue-Tek Optimum Cutting Temperature (OCT) compound (Sakura Finetek) and immediately frozen in 2-methylbutane precooled in liquid nitrogen. 5mm cryosections were mounted on gelatin coated 1.5 glass coverslips. Tissue sections were fixed in 4% paraformaldehyde, permeabilized with 0.2% Triton-X 100/PBS and blocked with 2% BSA/1% normal donkey serum/PBS prior to incubation with antibodies. The primary antibodies included: rabbit polyclonal anti-desmin (1:30) (ImmunoBioscience RP-4023-04), mouse monoclonal anti-sarcomeric a-actinin (1:1000) (Clone EA-53; Sigma A7811), and mouse monoclonal anti-desmoplakin 1/2 (1:1000) (Clone DP-2.15; AbDSerotec 2722-5204) antibodies. The secondary antibodies, obtained from Jackson Immunoresearch Laboratories, included: Alexa Fluor 488 goat anti-mouse IgG (1:500), Alexa Fluor 488 goat anti-rabbit IgG (1:500), Texas Red goat anti-mouse IgG (1:500), and Texas Red goat anti-rabbit IgG (1:500). Coverslips were mounted onto slides with Aqua Poly/Mount (Polysciences Inc.). All sections were analyzed on a Deltavision RT system with 100x (1.3 NA) objective and a CoolSnap HQ charge-coupled device camera (Photometrics) using softWoRx 3.5.1 software. Images were prepared for presentation using Photoshop CS (Adobe Systems).

### TUNEL analysis

Cell death on sections was detected using Roche In Situ cell death detection kit Fluorescein (11684795910).

### Yeast two-hybrid assay

A full-length BAF60c expression construct was used as a bait in a yeast two-hybrid assay conducted by Hybrigenics (http://www.hybrigenics-services.com/), using a human fetal/adult heart library.

### Luciferase Assay

The *Myl1* luciferase construct has been described (Creemers et al., 2006). Combined DNA vectors were transfected into early exponential stage 10T1/2 cells cultured in 6-well dishes with Fugene 6 (Roche, 1181443001) following the product manual. After culturing for another 40-48 hrs, the cells were lysed and luciferase activity analyzed with Dual-Luciferase Reporter Assay System (Promega E1910). The luciferase activity was normalized with renilla activity. Three biological replicates were prepared for each combination.

### GST-pulldown assay

^35^S labeled proteins (TBX5, NKX2-5, RBPjk, NICD, BAF60c serial deletions, Myocardin serial deletions (Wang et al., 2001)) were synthesized with the TnT SP6 coupled reticulocyte lysate system (Promega, L4600) or TnT T7 coupled reticulocyte lysate system (Promega L4610) and labeled with ^35^S Methionine (Perkin Elmer NEG709A). 5 μl of each synthesized protein was analyzed with SDS-PAGE gel and exposed to X-ray film for evaluation. GST-BAF60c, GST-RBPjk, GST-TBX5 and GST were expressed in *E.Coli* strain BL21 and purified with Glutathione Sepharose 4B (GE Healthcare, 17-0756-01). The Glutathione sepharose 4B beads were incubated with ^35^S labeled target proteins overnight at 4°C and washed with PBST for 3 times. The beads were then boiled in loading buffer. The protein was analyzed with SDS-PAGE gel followed by autoradiography

### Statistics

Data were expressed as mean±SEM. Differences among multiple experimental groups were evaluated by ANOVA followed by post-hoc Fisher’s LSD test. Pairwise comparisons were evaluated by student’s *t*-tests. *p* values < 0.05 were considered as significant.

## Acknowledgements

We thank the Toronto Center for Phenogenomics for chimera production, A. Williams and S. Thomas (Gladstone Bioinformatics Core) for data analysis, L. Ta (Gladstone Genomics Core) and J. Gorham (Harvard Medical School) for RNAseq library preparation, Yew Meng and Aina Tilups (Sickkids Pathlogy Dept) for electron microscopy, J.N. Wylie for luciferase assay, and G. Howard for editing.

